# Cell-type selective deletion of RSK2 reveals insights into altered signaling in Coffin-Lowry Syndrome

**DOI:** 10.1101/156257

**Authors:** Hu Zhu, Ryan T. Strachan, Daniel J. Urban, Martilias S. Farrell, Wesley K. Kroeze, Justin G. English, Reid H.J. Olsen, Bryan L. Roth

**Affiliations:** Department of Pharmacology, School of Medicine, University of North Carolina at Chapel Hill, Chapel Hill, NC; Carolina Institute for Developmental Disabilities, University of North Carolina, Chapel Hill, NC, USA; Program in Neuroscience, University of North Carolina, Chapel Hill, NC, USA; Lineberger Comprehensive Cancer Center, University of North Carolina, Chapel Hill, NC, USA; Division of Chemical Biology and Medicinal Chemistry, School of Pharmacy, University of North Carolina, Chapel Hill, NC, USA; NIMH Psychoactive Drug Screening Program, University of North Carolina, Chapel Hill, NC, USA

**Keywords:** 5-HT2A serotonin, Coffin-Lowry Syndrome, GPCR, Scaffolding proteins

## Abstract

Coffin-Lowry syndrome (CLS) is an X-linked syndromic form of mental retardation characterized by various skeletal dysmorphisms, moderate to severe mental retardation, and in some cases, psychosis. CLS is caused by loss-of-function mutations of the p90 ribosomal S6 kinase 2 (RPS6KA3) gene encoding a growth factor-regulated serine/threonine kinase, ribosomal S6 kinase 2 (RSK2). We previously identified RSK2 as a novel interacting protein that tonically inhibits 5-HT_2A_ receptor signaling by phosphorylating Ser-314 within the third intracellular loop. To determine if RSK2 inhibits 5-HT_2A_ receptor signaling *in vivo* and whether disruption of RSK2 could lead to schizophrenia-like behaviors - as is seen in some CLS patients - we genetically disrupted the function of RSK2 either globally or selectively in forebrain pyramidal neurons in mice. Both global and forebrain-selective RSK2 deletion augmented the locomotor responses to the psychotomimetic drugs phencyclidine (PCP) and amphetamine (AMPH). Significantly, forebrain-selective deletion of RSK2 augmented 5-HT_2A_ receptor signaling as exemplified by enhanced 5-HT_2A_-mediated c-fos activation and head-twitch response without altering the levels or distribution of 5-HT_2A_ receptor protein. Thus, RSK2 modulates 5HT_2A_ receptor function *in vivo,* and disruption of RSK2 leads to augmented psychostimulant-induced responses reminiscent of those seen in many animal models of schizophrenia. These findings strengthen the association between 5-HT_2A_ receptor dysfunction and psychosis, and provide a potential mechanism underlying the schizophrenia-like symptoms present in some CLS patients.

**Highlights:**

- Global and cell-type-specific RSK2 knock-out mice were assessed behaviorally and pharmacologically
- Augmentation of amphetamine and PCP locomotor responses were seen in both global and forebrain-specific RSK2 KO mice
- Augmentation of 5-HT2A serotonin receptor function but not number was also observed
- These alterations reveal insights into mechanisms potentially responsible for behavioral sequlae of Coffin-Lowry Syndrome

## 1. INTRODUCTION^1^

Coffin‒Lowry syndrome (CLS) is an X chromosome-linked mental retardation disorder characterized by growth defects, hypotonia, psychomotor retardation and distinct facial, hand and skeletal malformations (A. Hanauer and I. D. Young, 2002). Individuals with CLS frequently display the symptoms of autism spectrum disorder as well as psychosis (U. Sivagamasundari et al., 1994). CLS is caused by a variety of heterogeneous loss-of-function mutations in the ribosomal protein S6 kinase, 90kDa, polypeptide 3 (RPS6KA3) gene, which maps to Xp22.2 (E. Trivier et al., 1996). Over 150 distinct mutations have been discovered, with most being unique to individual affected families (J. Delaunoy et al., 2001; J. P. Delaunoy et al., 2006; P. M. Pereira et al., 2010; A. Schneider et al., 2013). The RPS6KA3 gene encodes the growth factor-regulated serine/threonine kinase, ribosomal S6 kinase 2 (RSK2). Mutations of the RPS6KA3 gene lead to premature termination of translation and/or to loss of phosphotransferase activity of RSK2 (A. Hanauer and I. D. Young, 2002; P. M. Pereira et al., 2010). Global RSK2 knockout (KO) mice display profound growth retardation, progressive bone loss, memory deficits, and cognitive deficiencies (S. D. Dufresne et al., 2001; X. Yang et al., 2004; R. Poirier et al., 2007).

RSK2 is a member of the ribosomal S6 kinase family which includes RSK1-RSK4. The RSK2 protein contains two functional kinase domains: the N-terminal kinase domain belongs to the AGC (protein kinase A, G and C) family, and the C-terminal kinase domain belongs to the CAMK (calcium/calmodulin-dependent protein kinase) family. RSK2 is a multifunctional effector of the mitogen-activated protein kinase (MAPK) signaling pathway (J. Xing et al., 1996) and is phosphorylated and activated by extracellular signal-regulated kinase (ERK) in response to growth factors, polypeptide hormones and neurotransmitters. Upon activation, RSK2 can phosphorylate a variety of substrates both in the cytosol (e.g., Glycogen synthase kinase 3 beta (GSK3β), L1 cell adhesion molecule (L1CAM), Eukaryotic translation initiation factor 4b (eIF4B), BCL2-associated agonist of cell death (BAD) and SH3 and multiple ankyrin repeat domains (SHANK)) and in the nucleus (e.g., cAMP response element-binding protein (CREB), Estrogen receptor (ER), Nuclear hormone receptor 77 (Nur77), FBJ murine osteosarcoma viral oncogene homolog (c-fos) and Activating transcription factor 4 (ATF4)) to regulate cell proliferation, differentiation and survival (Y. Romeo et al., 2012).

Loss-of-function of RSK2 in humans leads to Coffin-Lowry syndrome, which includes behavioral characteristics of a schizophrenia-like psychosis in some individuals (R. A. Collacott et al., 1987; U. Sivagamasundari et al., 1994; A. G. Hunter, 2002). However, it is unknown how RSK2 mutations lead to this psychosis-like condition. In previous studies (D. J. Sheffler et al., 2006), we discovered that RSK2 exerts a tonic brake on the signaling of a variety of G protein-coupled receptors (e.g., P2Y-purinergic, PAR-1-thrombinergic, β1-adrenergic, and bradykinin-B receptors). We also found that RSK2 directly interacts with the 5-HT_2A_ receptor and phosphorylates intracellular loop 3 at the conserved Ser-314 to attenuate 5-HT_2A_ receptor signaling (R. T. Strachan et al., 2009). Accordingly, we hypothesized that genetic deletion of the RSK2 gene may lead to an augmentation of 5-HT_2A_ receptor signaling and induction of a psychosis-like behavior in mice. To test this hypothesis, here we used two mouse models: a global RSK2 KO mouse and a newly created conditional RSK2 KO mouse. We found that both global and forebrain-specific deletion of RSK2 augments 5-HT_2A_ receptor signaling and some of the psychotomimetic actions of PCP and amphetamine.

## 2.0 MATERIALS AND METHODS

### 2.1 Animals

Mice were maintained in a 12-h light/dark cycle with unlimited access to water and laboratory chow. All procedures were conducted in strict compliance with the “Guide for the Care and Use of Laboratory Animals” (Institute of Laboratory Animal Resources, National Research Council, 1996 edition) and approved by the Institutional Animal Care and Use Committee of the University of North Carolina. RSK2 KO mice (C57BL/6J) were generated and provided by Dr. André Hanauer (Institute of Genetics, and Molecular and Cellular Biology, Illkirch, CEDEX, France) (R. Poirier et al., 2007).

### 2.2 Creation of a conditional RSK2 KO mouse

Briefly, a targeting vector for the mouse RSK2 gene was constructed. RSK2 exon 4 was flanked by loxP sites, and two positive selection markers flanked by flippase (Flp) recognition target sites FRT (NeoR, neomycin resistance) and F3 (PuroR, Puromycin resistance) were inserted into introns 3 and 4, respectively. The targeting vector was transfected into the TaconicArtemis C57BL/6N Tac ES cell line by Taconic Farms, and homologous recombinant clones were isolated using double positive selection markers. The targeted ES cells were injected into blastocysts to generate chimeras. Chimerism was measured by coat color contribution of ES cells to BALB/C host (black/white). Highly chimeric mice were bred with Flp-Deleter mice (Taconic Farms) to remove the selection markers. The conditional allele (RSK2^flox^) was confirmed by genotyping. The tail DNA was amplified by PCR with a primer pair directed against the third intron (forward primer, 5′- CCTTAAGTTACAACCCTAGCATCC -3′; reverse primer, 5′- TTCCTTTATTACAGCCAAGACTCC -3′), which generated a 525-bp amplicon in the conditional allele and a 353bp amplicon in the wild type allele. Mice carrying a conditional allele were then backcrossed to the C57 BL/6 mice for more than 10 generations.

To generate forebrain-selective RSK2 KO mice, homozygous male EMX1-Cre transgenic mice (Emx1^tm1(cre)Krj^, Jackson Labs) were crossed with heterozygous female RSK2^flox/+^ mice. All offspring will contain Cre recombinase transgene and since the RSK2 gene is located on the X chromosome, half of the male offspring will contain RSK2^flox^ allele and half will contain the WT allele. To detect the Cre recombinase transgene, PCR was carried out using the primers 5′-GCGGTCTGGCAGTAAAAACTATC-3′ and 5′-GTGAAACAGCATTGCTGTCACTT-3′, which generated a 102-bp amplicon from the Cre-coding region.

### 2.3 Quantitative reverse transcriptase-PCR

EMX-RSK2 KO and littermate control mice were injected with the 5-HT_2_-selective agonist 2,5-dimethoxy-4-iodoamphetamine (DOI, 0.5 mg/kg, i.p.) and after 1 h, mice were euthanized, and total RNA was immediately isolated from micro-dissected frontal cortex tissue using Trizol (Invitrogen, Carlsbad, CA, USA). 10 μg of RNA was treated with DNAse (DNA-free, Ambion, Austin, TX, USA), and first strand complementary DNA was synthesized from 2 μg of the DNase-treated RNA using the Superscript III RNase H Reverse Transcriptase kit (Invitrogen), with random hexamer primers (Invitrogen). Mouse c-Fos complementary DNA was amplified using Power SYBR Green PCR master mix (Applied Biosystems, Invitrogen, Carlsbad, CA, USA) in the 7500 Real-time PCR System (Applied Biosystems). The relative amount of c-Fos specific mRNA was determined by normalizing with β-actin mRNA present in the same samples and is expressed as fold above the saline-treated littermate control mice. The following gene-specific primers were used in PCR—mouse c-Fos: sense 5′-CAACACACAGGACTTTTGCG-3′, anti-sense 5′-GGAGATAGCTGCTCTACTTTG-3′; mouse β-actin: sense 5′-TGTTACCAACTGGGACGACA-3′; anti-sense 5′-CTGGGTCATCTTTTCACGGT-3′.

### 2.4 Western blot

Animals were sacrificed, and prefrontal cortex samples were harvested. The sample tissues were homogenized in ice-cold RIPA buffer (50mM Tris-Cl, pH 8.0; 150mM NaCl, 0.1% SDS, 1% NP-40, 0.5% Na-deoxycholate, plus protease inhibitor cocktail (50ml RIPA buffer/tablet, Roche)) and allowed to lyse on ice for 30 min. After centrifuging at 25,000 × g for 30min at 4°C, supernatants were transferred to a new tube. 20ug total protein were loaded and separated by SDS-PAGE gel, transferred to PVDF membranes, and probed with rabbit monoclonal anti-RSK2 antibody (1:1,000, Cell Signaling) at 4°C overnight. After three washes, the blot was probed with horseradish peroxidase-conjugated goat anti-rabbit IgG antibody (1:2,000, life technology) at room temperature for 1 hr. The blot was washed three times and incubated with chemiluminescent substrate (Thermo Scientific) and imaged in a Kodak Imager. After imaging, the blot was extensively washed and re-probed with mouse monoclonal anti-β-actin antibody (1:10,000, Sigma).

### 2.5 Immunohistochemistry

Immunohistochemistry was performed as previously described (A. I. Abbas et al., 2009). Briefly, adult mice were anesthetized and perfused with 4% parafromaldehyde (PFA) in 1xPBS. Mouse brains were harvested, postfixed in 1xPBS/4% PFA overnight, and then dehydrated in 1xPBS /30% sucrose until they sank. 40μm brain sections were cut and mounted onto poly-l-lysine coated microscope slides. To counterstain the neurons, NeuroTrace 530/615 (1:100, Invitrogen) was applied to slides and incubated at room temperature for 30min. After extensive washes, the sections were permeabilized with 1xPBS/0.4% Triton X-100 for 1hr, and then blocked with 1xPBS/0.4% Triton X-100 solution containing 1% bovine serum albumin and 5% normal goat serum at room temperature for 1hr. Primary antibodies diluted in the blocking solution were then applied to the slides and incubated at 4°C overnight. The following primary antibodies were used: mouse monoclonal anti-RSK2 antibody (1:200, Abcam) and rabbit polyclonal anti-5HT2A antibody (1:200, Neuromics). Following the wash with 1xPBS/0.4% Triton X-100, slides were incubated with either AlexaFluor 488 goat anti-mouse antibody or AlexaFluor 594 goat anti-rabbit antibody (1:250, Invitrogen) diluted in the blocking solution at room temperature for 1hr. Fluorescent images were collected on a Nikon 80i Research Upright Microscope (Nikon, Tokyo, Japan) equipped with Surveyor Software with TurboScan (Objective Imaging, Kansasville, WI). Tiled images were collected with a Qimaging Retiga-EXi camera (Qimaging, Surrey, BC, Canada). The signal intensity was quantified using Image J software (NIH). Raw images were corrected by subtracting the average intensity value of the background. Three regions of interest (ROIs), 200 by 200 pixels, were randomly selected for layer V and peripheral layers; the mean intensity was then measured for each ROI, and these were averaged for each brain section.

### 2.6 Head-twitch response

Head-twitch response was tested as reported previously (J. A. Allen et al., 2011). Briefly, animals were transferred to the procedure room and acclimated to the new environment for at least 30min. Animals were injected intraperitoneally (i.p.) with various doses of DOI or vehicle and placed into the center of a Plexiglas cage for 20 min, during which time head-twitch behavior was scored (5 min bins) by an experienced observer blind to genotype and drug treatment. Head-twitch behavior is defined as a rapid rotational jerk of the head, which is different from grooming or scratching behaviors (J. Gonzalez-Maeso et al., 2007).

### 2.7 Open field test

Exploration in a novel environment was assessed by a two hour trial in an open field (40 cm x 40 cm x 30 cm) crossed by a grid of photobeams (VersaMax system, AccuScan Instruments). Counts were taken of the number of photobeams broken during the trial in five-minute intervals, with separate measures for horizontal activity (total distance traveled), stereotypy activity (repeated breaking of the same set of photobeams), and vertical activity (rearing movements). For pharmacological studies, animals were injected IP with either MDL 100907 (0.2 or 0.5 mg/kg) or vehicle (0.9% saline) and placed into the activity box. After 30min, animals were injected IP with PCP or AMPH (6 mg/kg) and immediately returned to the activity box for 90min. Activity was monitored during whole period.

### 2.8 Elevated plus-maze test

Mice were given one 5-min trial on a metal plus-maze, which had two closed arms with walls 20 cm in height, and two open arms. The maze was elevated 50 cm from the floor, and the arms were 30 cm long. Animals were placed on the center section (8 cm x 8 cm), and allowed to freely explore the maze. Arm entries were defined as all four paws entering an arm. Entries and time in each arm were recorded during the trial by a blinded human observer via computer coding.

### 2.9 Accelerating rotarod

Subjects were tested for motor coordination and learning on an accelerating rotarod (Ugo Basile, Stoelting Co., Wood Dale, IL). For the first test session, animals were given three trials, with 45 seconds between each trial. Two additional trials were given 48 hours later. Rpm (revolutions per minute) was set at an initial value of 3, with a progressive increase to a maximum of 30 rpm across five minutes (the maximum trial length). Measures were taken for latency to fall from the top of the rotating barrel in five minute test periods.

### 2.10 Social approach test

The three-chamber social approach test was designed to assess the sociability and the preference of social novelty of animals. All animal behaviors were monitored and analyzed by an automated tracking system (Noldus Ethovision). The test subject was first placed in the middle chamber and allowed to explore for ten minutes. After the habituation period, an unfamiliar C57BL/6J mouse (stranger 1) was placed in one of the side chambers enclosed in a small wire cage. An identical empty wire cage was placed in the opposite side of the chamber. The subject was then allowed to explore three chambers for a ten-minute session and the amount of time subject spent exploring the unfamiliar mouse or empty cage was recorded. At the end of the sociability test, each mouse was further tested for exploration of a new unfamiliar mouse (stranger 2) placed in the previously empty wire cage. Under these conditions the test mouse had a choice between the first, already-investigated mouse (stranger 1) and the novel unfamiliar mouse (stranger 2). The same measures were taken as with the sociability test.

### 2.11 Morris water maze

The Morris water maze task was described in our previous study (H. Zhu et al., 2014). Briefly, mice were given a visual cue test first to evaluate the visuo-motor abilities and the motivation of the animals. After the visual cue test, mice were trained in the hidden platform test as described previously. Each animal was given 4 trials per day, across 6 days, to learn the location of the submerged platform. The criterion for learning was an average latency of 15 sec or less to locate the hidden platform. After meeting this criterion, mice were then tested for spatial memory during a 1min trial in the pool with the platform removed. Spatial memory was evaluated by measuring the percent of time mice spent in the quadrant where the hidden platform was located.

### 2.12 Fear conditioning

Conditioned fear was evaluated using a Near-Infrared Video Fear Conditioning system (MED Associates, St. Albans, VT). In the training phase, mice received 3 pairings of a 30-sec, 90 dB, 5 kHz tone (conditioned stimulus, CS) and a 2-sec, 0.6 mA foot shock (unconditioned stimulus, US). Contextual memory was evaluated in the original training chamber 24 hours following the training phase, whereas cued memory was evaluated in a new chamber 48 hours following the training phase. Periods of freezing (no movement for 0.5 sec) were automatically measured by the image tracking software (Med Associates, St Albans, VT).

### 2.13 Prepulse inhibition of startle response

The acoustic startle test was used to evaluate reactivity to an auditory stimulus and sensorimotor gating as previously described (J. A. Allen et al., 2011). Mice were tested with a San Diego Instruments SR-Lab system. Briefly, mice were placed in a small Plexiglas cylinder within a larger, sound-attenuating chamber. The cylinder was seated upon a piezoelectric transducer, which allowed vibrations to be quantified and displayed on a computer. The chamber included a ceiling light, fan, and a loudspeaker for the acoustic stimuli. Background sound levels (70 dB) and calibration of the acoustic stimuli were confirmed with a digital sound level meter (San Diego Instruments). Each session consisted of 42 trials that began with a five-minute habituation period. There were 7 different types of trials: no-stimulus trials, trials with the acoustic startle stimulus (40 msec; 120 dB) alone, and trials in which a prepulse stimulus (20 msec, either 74, 78 or 82dB) occurred 100 ms before the onset of the startle stimulus. Measurements were taken of the startle amplitude for each trial across a 65-msec sampling window, and an overall analysis was performed for each subject’s data for levels of prepulse inhibition at each prepulse sound level (calculated as 100 - [(response amplitude for prepulse stimulus and startle stimulus together / response amplitude for startle stimulus alone) x 100]. For pharmacological studies, animals were injected IP with vehicle (0.9% saline) or MDL 100907 (0.2 or 0.5 mg/kg) and returned to their home cage. After 30 min, animals were injected IP with phencyclidine (4 or 6 mg/kg) and immediately placed into the PPI apparatus.

### 2.14 Receptor autoradiography

5-HT_2A_ receptor autoradiography was performed as previous reported (J. A. Allen et al., 2011). Briefly, brains were quickly removed, frozen on dry ice, and stored at -80°C until processing. Coronal brain sections (20 μm) (Bregma: 2.10 mm to Bregma 1.54 mm) were cut on a cryostat, thaw-mounted onto slides (Fisher Scientific Tissue Path Superfrost Plus Gold Slides, #15-188-48), and vacuum desiccated at 4°C overnight. Sections were incubated with ^125^I-DOI ((+/-)-1-(2,5-dimethoxy-4-iodophenyl)-2-aminopropane HCL) (200 pM, 5-HT_2A/2C_ receptor agonist) for 1 hour at RT in standard binding buffer (50mM TrisHCl, pH 7.4; 10 mM MgCl, 0.1mM EDTA) to determine total binding. Ritanserin (1 uM; 5-HT_2A_/5-HT_2C_ receptor antagonist) was added to the incubation mixture to determine non-specific binding. Sections were washed (3×10 min) in ice-cold binding buffer (50 mM TrisHCl, pH 7.4), rinsed in ice-cold water (to remove residual salts), then air-dried and exposed to hyperfilm (GE Healthscience) for one week. Films were developed and photographed with a digital camera system (Metaview). Image analysis was performed using ImageJ software to define regions of interest (ROI) in cortical layers 5 and 6 and this ROI was transposed onto the non-specific binding images. Average pixel intensities of the nonspecific ROI was subtracted from the total ROI to obtain qualitative specific binding data.

### 2.15 Statistical Analysis

For quantitation of immunoblots, immunostaining, mRNA expression and autoradiography, or other two-group comparisons, two-tailed unpaired t-tests were used to compare whether there is a significant difference between two groups. All behavioral data were analyzed by two-way ANOVA followed by Bonferroni post tests for comparing multiple groups. Comparisons were considered significant if the p-value was <0.05.

## 3.0 RESULTS

### 3.1 Deletion of RSK2 augments psychotomimetic drug actions

Deleting RSK2 in mice leads to a recapitulation of many of the symptoms of CLS patients, including deficits in body weight, adipose mass, glycogen metabolism, skeletal development, and learning and memory (S. D. Dufresne et al., 2001; K. El-Haschimi et al., 2003; X. Yang et al., 2004; R. Poirier et al., 2007). As some individuals with CLS exhibit a schizophrenia-like psychosis (R. A. Collacott et al., 1987; U. Sivagamasundari et al., 1994; A. G. Hunter, 2002), we evaluated two well-validated behavioral models that have been extensively used in various mouse models of psychosis and in testing antipsychotic drugs in mice: locomotion in the open field and prepulse inhibition of startle response (PPI) (S. B. Powell et al., 2009) after administration of the psychostimulants PCP and AMPH (D. C. Javitt and S. R. Zukin, 1991). Consistent with our previous work (A. I. Abbas et al., 2009; J. A. Allen et al., 2011), administration of 6mg/kg PCP (Figure 1A) or AMPH (Figure 1C) induced hyperlocomotion in WT mice. Intriguingly, the hyperlocomotion induced by PCP (interaction of genotype and time, F(17,459)=2.35, p=0.0018; effect of genotype, F(1,459)=4.737, p=0.0384, two-way ANOVA) or AMPH (interaction of genotype and time, F(17,238)=3.899, p<0.0001; effect of genotype, F(1,238)=7.257, p=0.0175, two-way ANOVA) was significantly greater in global RSK2 KO mice compared with WT mice (Figure 1A and 1C). The selective 5-HT_2A_ receptor antagonists MDL 100,907 (MDL) (P. N. Yadav et al., 2010; J. A. Allen et al., 2011) and pimavanserin (P. N. Yadav et al., 2011; H. Y. Meltzer and B. L. Roth, 2013) have previously been demonstrated to attenuate PCP-induced hyperlocomotion in mice. As shown in Figure 1B, pretreatment with 0.5mg/kg MDL significantly decreased PCP-induced locomotion in both genotypes. However, MDL was less effective in normalization of PCP-induced hyperlocomotion in RSK2 KO mice compared with WT mice at time point 35 minute and 40 minute (interaction of genotype and time, F(17,476)=2.913, p<0.0001; effect of genotype, F(1,476)=3.582, p=0.0688, two-way ANOVA). Pretreatment with 0.5 mg/kg MDL also decreased AMPH-induced locomotion in both genotypes. However, as was observed with PCP (Figure 1B), MDL was less effective in global RSK2 KO mice compared with WT mice (interaction of genotype and time, F(17,238)=3.43, p<0.0001; effect of genotype, F(1,238)=7.155, p=0.0181, two-way ANOVA) (Figure 1D).

**Figure 1.**
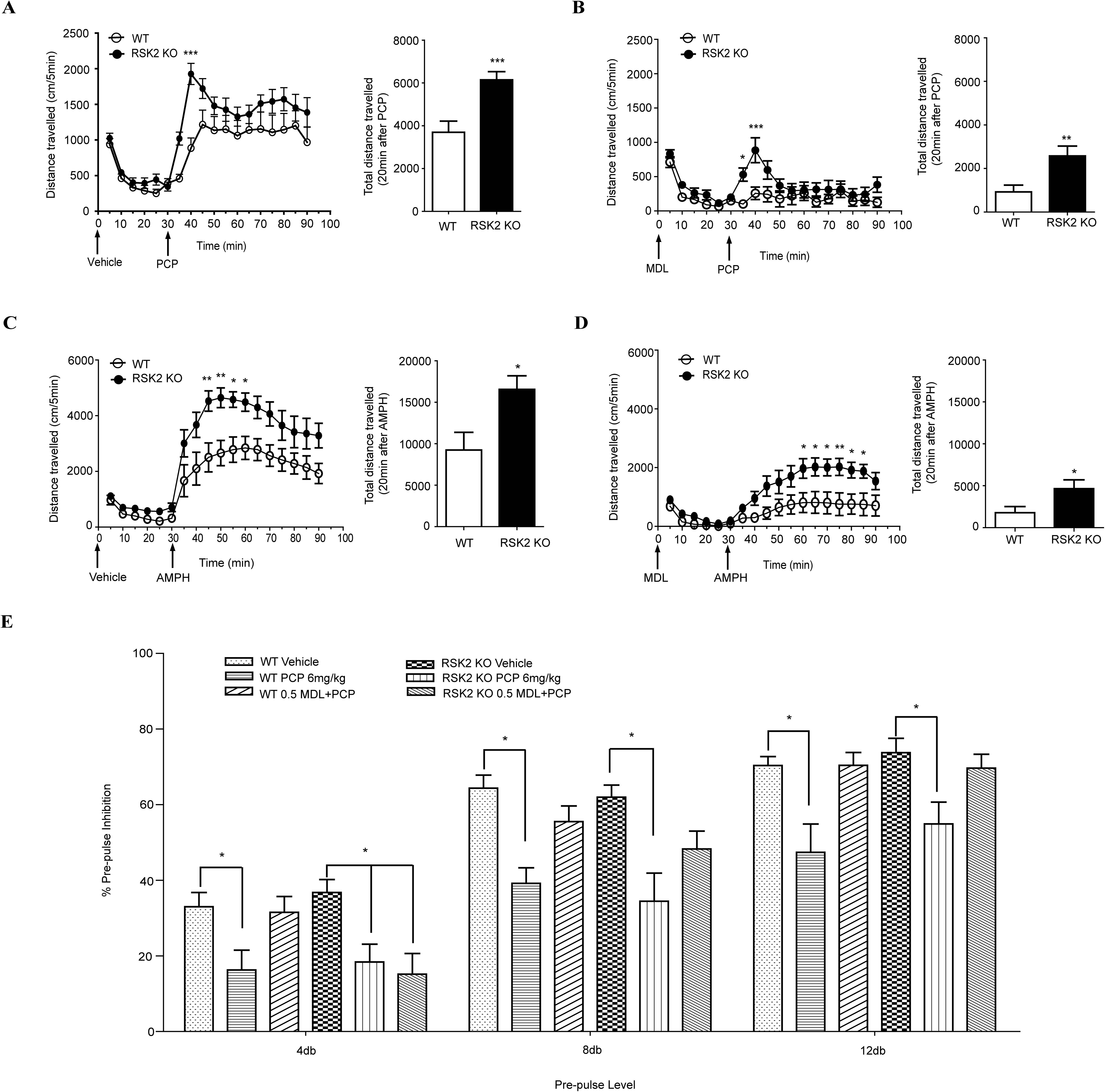
Disruption of the RSK2 gene attenuated the effectiveness of 5-HT_2A_ receptor antagonist on normalizing the PCP or AMPH-induced psychosis. **A**. Disruption of RSK2 gene significantly increased PCP-induced hyperlocomotion (interaction of genotype and time, F(17,459)=2.35, p=0.0018; effect of genotype, F(1,459)=4.737, p=0.0384, two-way ANOVA). Animals were injected with vehicle and placed into open field chambers, 30 min later mice were injected with 6mg/kg PCP and the total distance traveled was determined. **B**. Disruption of RSK2 gene significantly decreased the effectiveness of MDL on normalizing the PCP-induced locomotion (interaction of genotype and time, F(17,476)=2.913, p<0.0001; effect of genotype, F(1,476)=3.582, p=0.0688, two-way ANOVA). Animals were pretreated with 0.5mg/kg MDL and placed into open field chambers, 30 min later mice were injected with 6mg/kg PCP and the total distance traveled was determined. **C**. Disruption of RSK2 gene significantly increased AMPH-induced hyperlocomotion (interaction of genotype and time, F(17,238)=3.899, p<0.0001; effect of genotype, F(1,238)=7.257, p=0.0175, two-way ANOVA). Animals were injected with vehicle and placed into open field chambers, 30 min later mice were injected with 6mg/kg AMPH and the total distance traveled was determined. **D**. Disruption of RSK2 gene significantly decreased the effectiveness of MDL on normalizing the AMPH-induced locomotion (interaction of genotype and time, F(17,238)=3.43, p<0.0001; effect of genotype, F(1,238)=7.155, p=0.0181, two-way ANOVA). Animals were pretreated with 0.5mg/kg MDL and placed into open field chambers, 30 min later mice were injected with 6mg/kg AMPH and the total distance traveled was determined. **E.** Disruption of RSK2 gene decreased the effectiveness of MDL on normalizing the PCP-induced disruption of sensorimotor gating. Animal received vehicle, 6mg/kg PCP, 0.5mg/kg MDL + 6mg/kg PCP. MDL significantly normalized the PCP-induced disruption of PPI in WT mice (F(1,40)=12.49, p=0.0021, two-way ANOVA), but not in RSK2 KO mice (F(1,42)=0.5965, p=0.4485, two-way ANOVA, n=11). Two-way ANOVA followed by Bonferroni post test for multiple comparisons, * p<0.05, ** p<0.01, *** p<0.001.

We next examined the effect of global RSK2 deletion on PCP-induced disruption of PPI (i.e., an inhibitory process wherein a preceding weak stimulus inhibits the response to a closely following strong stimulus), which is a well-established mouse model of schizophrenia-like circuitry dysfunction (D. L. Braff et al., 2001; G. S. Linn and D. C. Javitt, 2001). As shown in Figure 1E, baseline levels of PPI were comparable between WT and global RSK2 KO mice. PCP significantly disrupted PPI in both genotypes at all prepulse levels. Similar to locomotion studies (Figure 1B), we observed that MDL was effective at normalizing PCP-induced disruption of PPI in WT mice (F(1,40)=12.49, p=0.0021, two-way ANOVA), but not in RSK2 KO mice (F(1,42)=0.5965, p=0.4485, two-way ANOVA) particularly at 4 and 8 db prepulse levels (Figure 1E). Taken together, locomotion and PPI studies support an enhancement of 5-HT_2A_-dependent signaling in global RSK2 KO mice.

### 3.2 Generation of forebrain-specific RSK2 KO mice

Given that the global RSK2 KO mouse has many non-neuronal phenotypes due to global RSK2 deletion, we undertook a more targeted approach by generating forebrain-specific conditional RSK2 KO mice. In these mice, exon 4 of the RSK2 gene was flanked by two LoxP sites that are removed by Cre recombinase, leading to a frameshift and premature termination of translation in exon 5. The resulting truncated protein contains only the first 81 amino acids of the RSK2 protein and lacks the kinase domain (Figure 2A). To selectively disrupt the RSK2 gene in the forebrain, we crossed heterozygous female conditional RSK2 mice (RSK2^flox/+^) with homozygous male EMX1-Cre mice (EMX^Cre/Cre^) selectively expressing Cre recombinase in the cortex, hippocampus and olfactory bulb (T. Iwasato et al., 2000). As the RSK2 gene is X-linked, male offspring were either EMX^Cre/+^RSK2^+^ or EMX^Cre/+^RSK2^flox^ (Figure 2B), with the EMX^Cre/+^RSK2^flox^ male mice possessing forebrain disruption of the RSK2 gene (hereafter referred to as EMX-RSK2 KO mice). Littermate EMX^Cre/+^RSK2^+^ male mice served as controls in all studies.

**Figure 2.**
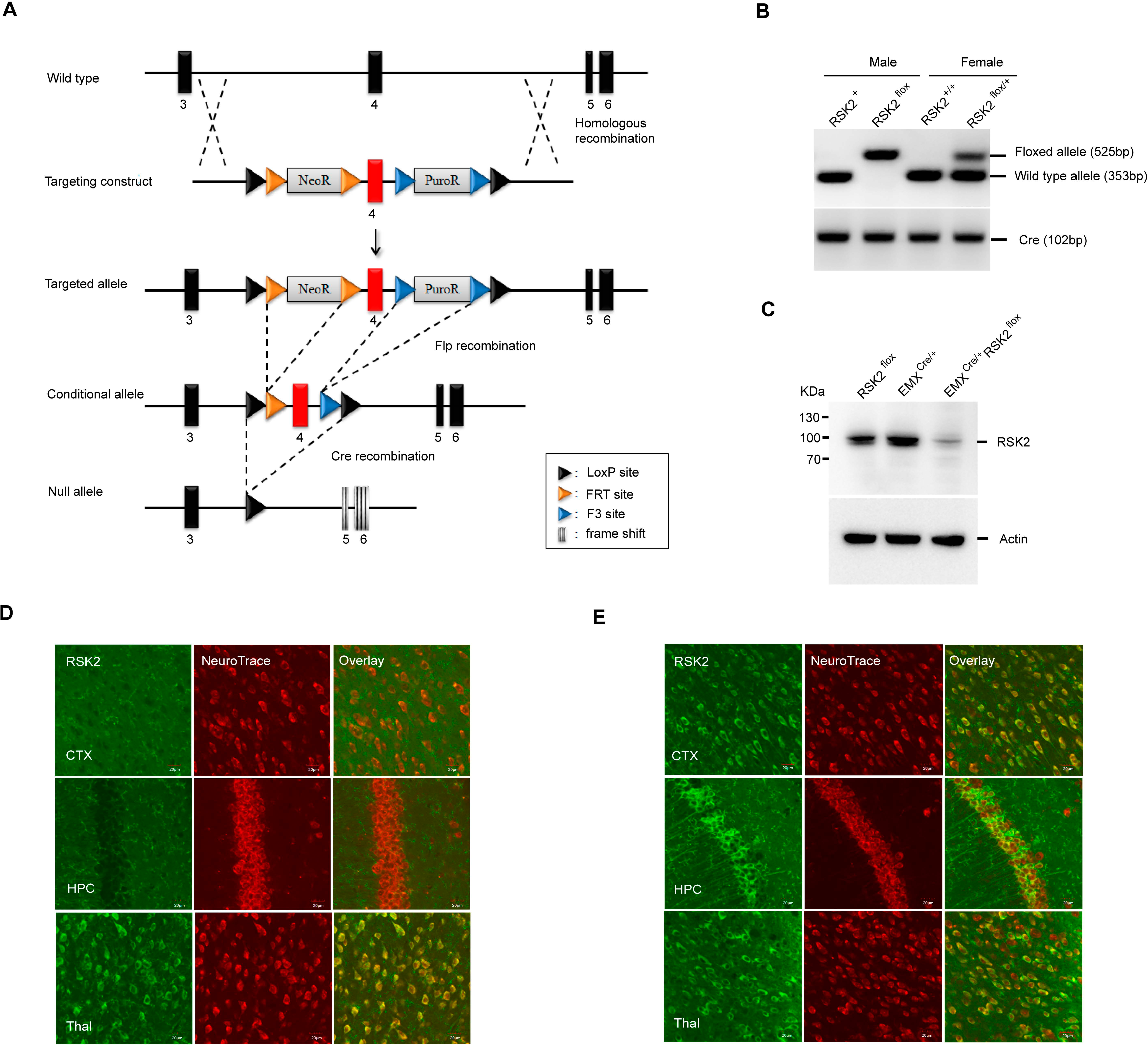
Forebrain-specific disruption of RSK2 gene in EMX-RSK2 KO mice. **A**. Schematic representation of the wild-type RSK2 allele, targeting construct, targeted allele, conditional allele, and null allele. Exon 4 is shown as a red box, and exons 3, 5, 6 are shown as black boxes. LoxP sequences are shown as black triangles, FRT sequences are shown as orange triangles, and F3 sequences are shown as blue triangles. Grey boxes indicate the neomycin (NeoR) and puromycin (PuroR) resistance sequences used for double positive selection. Striped boxes represent the frame shift of exons 5 and 6 after removing exon 4. The conditional KO allele was generated by partial excision of the targeted allele using Flp-Deleter mice, and the null allele was generated by excision of exon 4 from conditional allele using EMX-Cre mice. **B**. Genotyping of EMX-RSK2 KO mice. Male homozygous EMX-Cre mice were crossed with female heterozygous conditional RSK2 mice. Male litters showed either a 353bp wild type allele (lane 1) or 525bp conditional allele (lane 2). Female litters showed either wild type alleles (lane 3) or heterozygous alleles (wild type and conditional allele, lane 4). All litters showed a 102bp amplicon of EMX-Cre transgene in the low panel. **C**. Western blot analysis of RSK2 (~90 kDa) in protein lysates prepared from the cerebral cortex of conditional RSK2 mice (lane1), EMX-Cre mice (lane2) and EMX-RSK2 KO mice (lane3) at postnatal day 56. Comparable staining of β-actin was used to verify equivalent protein loading. **D, E**. Immunofluorescence staining analysis of RSK2 protein (green) in the cortex (CTX), hippocampus (HFC) and thalamus (Thal) from EMX-RSK2 KO mice (**D**) or littermate control mice (**E**). Neurons were counterstained with NeuroTace 530/615 (red). Scale bar is 20 um.

To confirm forebrain-specific disruption of RSK2, we analyzed RSK2 protein expression using Western blot and immunofluorescence microscopy. Western blot results showed that there was a single ~90 kDa band of RSK2 protein in conditional RSK2 mice and EMX^Cre/+^RSK2^+^ control mice, but only a weak band was present in the EMX-RSK2 KO mice (Figure 2C). Immunohistochemistry with the neuron-specific stain NeuroTrace showed that RSK2 was not expressed in cortical (CTX) and hippocampal (HPC) neurons of EMX-RSK2 KO mice (Figure 2D), although expression was retained in thalamic neurons (Thal) lacking Cre recombinase (Figure 2D). In contrast, RSK2 was expressed in the neurons of all three brain regions in control mice (Figure 2E). Taken together, we generated EMX-RSK2 KO mice with forebrain-specific disruption of RSK2.

### 3.3 EMX-RSK2 KO mice exhibit decreased sensitivity to 5-HT_2A_ antagonism

To determine whether forebrain-specific deletion of RSK2 leads to any intrinsic basal behavioral changes, animals were subjected to a battery of neurobehavioral tests in their naïve state. EMX-RSK2 KO mice showed normal growth, body weight, and mating behavior, unlike global RSK2 KO mice that have been reported to display decreases in growth and weight gain (X. Yang et al., 2004). Additionally, basal locomotion, anxiety, motor coordination, sociability, and learning and memory were comparable between the KO mice and littermate control mice, as demonstrated in open-field, elevated plus maze, accelerating rotarod, social approach, fear conditioning and Morris water maze (Figure 3A to 3G). Together, these data show no baseline behavioral changes after disruption of RKS2 in the forebrain, distinct from results obtained by others with global RSK2 KO mice.

**Figure 3.**
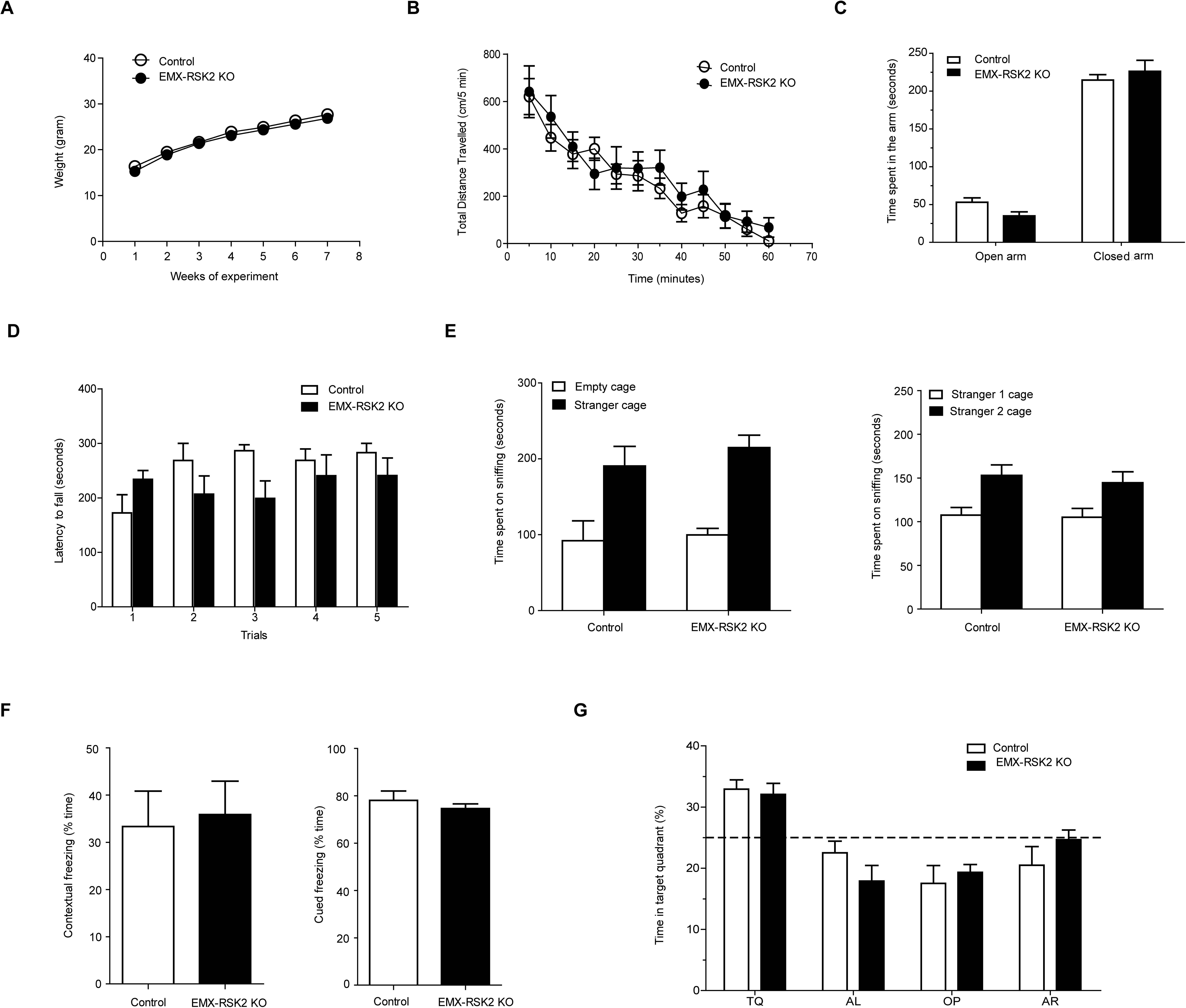
EMX-RSK2 KO mice exhibited normal growth and basal behaviors in their naïve state. EMX-RSK2 KO mice (n=7) showed a normal growth compared with control mice (n=6) (**A**, F(1,66)=1.253, p=0.2868, two-way ANOVA). In novelty induced locomotion test, there is a compatible locomotor response between two genotypes (**B**, F(1,121)=0.3097, p=0.589, two-way ANOVA). EMX-RSK2 KO mice showed a normal anxiety level in the elevated plus maze test (**C**, F(1,36)=0.4865, p=0.4944, two-way ANOVA) and a normal motor coordination in the accelerating rotarod test (**D**, F(1,44)=1.178, p=0.3009, two-way ANOVA). In the sociability and social novelty test, EMX-RSK2 KO mice showed a comparable preference compared with the littermate control mice (**E,** sociability, F(1,10),=2.167, p=0.1718; social novelty, F(1,11)=0.1802, p=0.6794, two-way ANOVA). EMX-RSK2 KO mice also showed a normal contextual and cued freezing behaviors in fear conditioning test (**F**, contextual freezing, t(11)=0.2441, p=0.8116; cued freezing, t(1,11)=0.8154, p=04322, unpaired t-test). In the Morris water maze test, EMX-RSK2 KO mice showed a comparable preference to the target quadrant during the memory test (**G**, F(1,33)=0.1732, p=0.6853). * p<0.05, ** p<0.01, *** p<0.001.

We next determined whether forebrain-specific RSK2 KO mice exhibit an augmented response to the psychotomimetic drugs PCP and AMPH. As seen in global RSK2 KO mice (Figure 1), PCP or AMPH administration induced hyperlocomotion in both control and EMX-RSK2 KO mice (Figure 4A and C). We observed no significant difference in PCP-induced hyperlocomotion between genotypes (interaction of genotype and time, F(17,340)=0.9219, p=0.5482; effect of genotype, F(1,340)=0.4912, p=0.4915, two-way ANOVA) (Figure 4A); however, we observed a significant increase in AMPH-induced hyperlocomotion in EMX-RSK2 KO mice (interaction of genotype and time, F(17,187)=5.725, p<0.0001; effect of genotype, F(1,187)=11.63, p=0.0058, two-way ANOVA) (Figure 4C). Next, we evaluated whether the PCP- or AMPH-induced hyperlocomotion in EMX-RSK2 KO mice could be effectively normalized by MDL. Injection of 0.2 mg/kg MDL 30min before PCP or AMPH administration significantly decreased PCP- and AMPH-induced locomotion in both genotypes (Figure 4B and D). However, as with the global RSK2 KO mice, MDL was less effective in EMX-RSK2 KO mice compared to littermate control mice (PCP- interaction of genotype and time, F(17,340)=4.283, p<0.0001; effect of genotype, F(1,340)=5.379, p=0.0311, two-way ANOVA) (AMPH - interaction of genotype and time, F(17,170)=4.591, p<0.0001; effect of genotype, F(1,170)=10.09, p=0.0099, two-way ANOVA) (Figure 4B and 4D).

**Figure 4.**
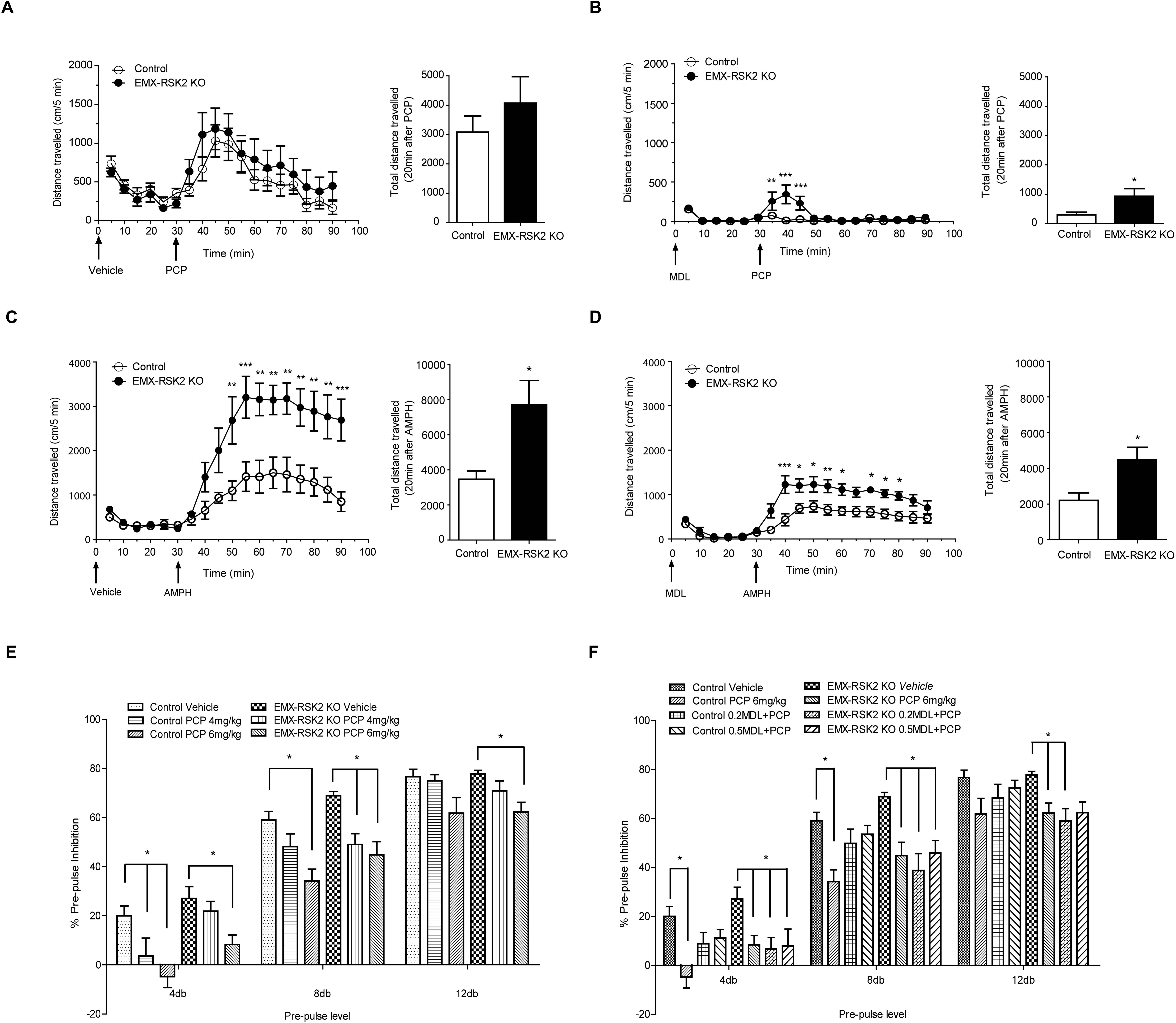
Forebrain-specific disruption of RSK2 gene attenuated the effectiveness of 5-HT_2A_ receptor antagonist on normalizing the PCP- or AMPH-induced psychosis. **A**. There was no significant difference between EMX-RSK2 KO mice and control mice in PCP-induced hyperlocomotion (interaction of genotype and time, F(17,340)=0.9219, p=0.5482; effect of genotype, F(1,340)=0.4912, p=0.04915, two-way ANOVA). Animals were injected with vehicle and placed into open field chambers, 30 min later mice were injected with 6mg/kg PCP and the total distance traveled was determined. **B**. Forebrain-specific disruption of RSK2 gene significantly decrease the effectiveness of MDL on normalizing the PCP-induced locomotion (interaction of genotype and time, F(17,340)=4.283, p<0.0001; effect of genotype, F(1,340)=5.379, p=0.0311, two-way ANOVA). Animals were pretreated with 0.2mg/kg MDL and placed into open field chambers, 30 min later mice were injected with 6mg/kg PCP and the total distance traveled was determined. **C**. Forebrain-specific disruption of RSK2 gene significantly increased AMPH-induced hyperlocomotion (interaction of genotype and time, F(17,187)=5.725, p<0.0001; effect of genotype, F(1,187)=11.63, p=0.0058, two-way ANOVA). Animals were injected with vehicle and placed into open field chambers, 30 min later mice were injected with 3mg/kg AMPH and the total distance traveled was determined. **D.**MDL was less effectiveness to normalize AMPH-induced hyperlocomotion (interaction of genotype and time, F(17,170)=4.591, p<0.0001; effect of genotype, F(1,170)=10.09, p=0.0099, two-way ANOVA). Animals were injected with MDL (0.2mg/kg) and placed into open field chambers, 30 min later mice were injected with 3mg/kg AMPH and the total distance traveled was determined. **E**. PCP disrupted the sensorimotor gating in both genotypes in a comparable manner. Animal received vehicle, 4mg/kg or 6 mg/kg PCP, and then followed by measuring the acoustic startle response. PCP significantly disrupted prespulse inhibition in control mice (F(2,64)=7.503, p=0.0021, two-way ANOVA; n=12) and EMX-RSK2 KO mice (F(2,82)=10.48, p=0.0002, two-way ANOVA; n=15). **F**. Forebrain-specific disruption of RSK2 gene significantly decreased the effectiveness of MDL on normalizing the PCP-induced disruption of sensorimotor gating. Animal received vehicle, 6mg/kg PCP, 0.2mg/kg MDL + 6mg/kg PCP or 0.5mg/kg MDL + 6mg/kg PCP. MDL normalized the disruption of PPI by PCP in control mice (F(2,66)=4.818, p=0.0146, two-way ANOVA; n=12), but it has no effect in EMX-RSK2 KO mice (F(2,84)=0.297, p=0.7446, twoway ANOVA; n=15). * p<0.05, ** p<0.01, *** p<0.001.

We next determined if forebrain-specific deletion of RSK2 decreased the ability of the 5-HT_2A_ antagonist MDL to normalize PCP-induced disruption of PPI as seen with the global RSK2 KO mice. As shown in Figure 4E, EMX-RSK2 KO mice exhibited both baseline and PCP-induced changes in PPI that were comparable to control mice. Injection of MDL (0.2 or 0.5mg/kg) 30min before PCP (6mg/kg) administration normalized PCP-induced disruption of PPI in control mice (F(2,66)=4.818, p=0.0146, two-way ANOVA); whereas it was not effective in EMX-RSK2 KO mice (F(2,84)=0.297, p=0.7446, two-way ANOVA) (Figure 4F). Similar to our observations with global RSK2 KO mice (Figure 1E), 5-HT_2A_ blockade was less effective at normalizing the psychotomimetic effects of PCP.

### 3.4 5-HT_2A_ signaling is augmented in EMX-RSK2 KO mice

Our previous studies (D. J. Sheffler et al., 2006; R. T. Strachan et al., 2009; R. T. Strachan et al., 2010a; R. T. Strachan et al., 2010b) demonstrated that RSK2 exerts a tonic brake on 5-HT_2A_ signaling pathway in fibroblasts *in vitro*, though no data was available regarding the actions in neurons. To test this *in vivo* in forebrain neurons, we evaluated neuronal signaling and behavioral activity in EMX-RSK2 KO mice following administration of the 5-HT_2A/2C_ receptor agonist DOI. DOI has been shown to selectively induce the expression of the immediate early response gene c-fos in the mouse cortex (J. L. Scruggs et al., 2000; A. I. Abbas et al., 2009), which is a marker of neuronal activation. Following 1 mg/kg DOI administration, both genotypes showed DOI-induced c-fos activation in the somatosensory cortex (SSC, Figure 5A) as demonstrated by immunohistochemistry with anti-c-fos antibody (Figure 5B). Notably, DOI-induced c-fos activation in EMX-RSK2 KO mice was significantly increased compared to littermate control mice (interaction of genotype and drug, F(1,24)=5.993, p=0.022, effect of genotype, F(1,24)=6.013, p=0.0218, two-way ANOVA) (Figure 5C). The augmented DOI-induced c-fos activation was also confirmed by analyzing c-fos gene expression via quantitative polymerase chain reaction (qPCR). We observed an increase of DOI-induced c-fos activation in both genotypes, with the level of c-fos mRNA induced by DOI in EMX-RSK2 KO mice significantly greater than in control mice (interaction of genotype and drug, F(1,32)=4.922, p=0.0337, effect of genotype, F(1,32)=4.538, p=0.0409, two-way ANOVA) (Figure 5D). Thus, forebrain-specific disruption of the RSK2 gene augments 5-HT_2A_ signaling *in vivo*.

**Figure 5.**
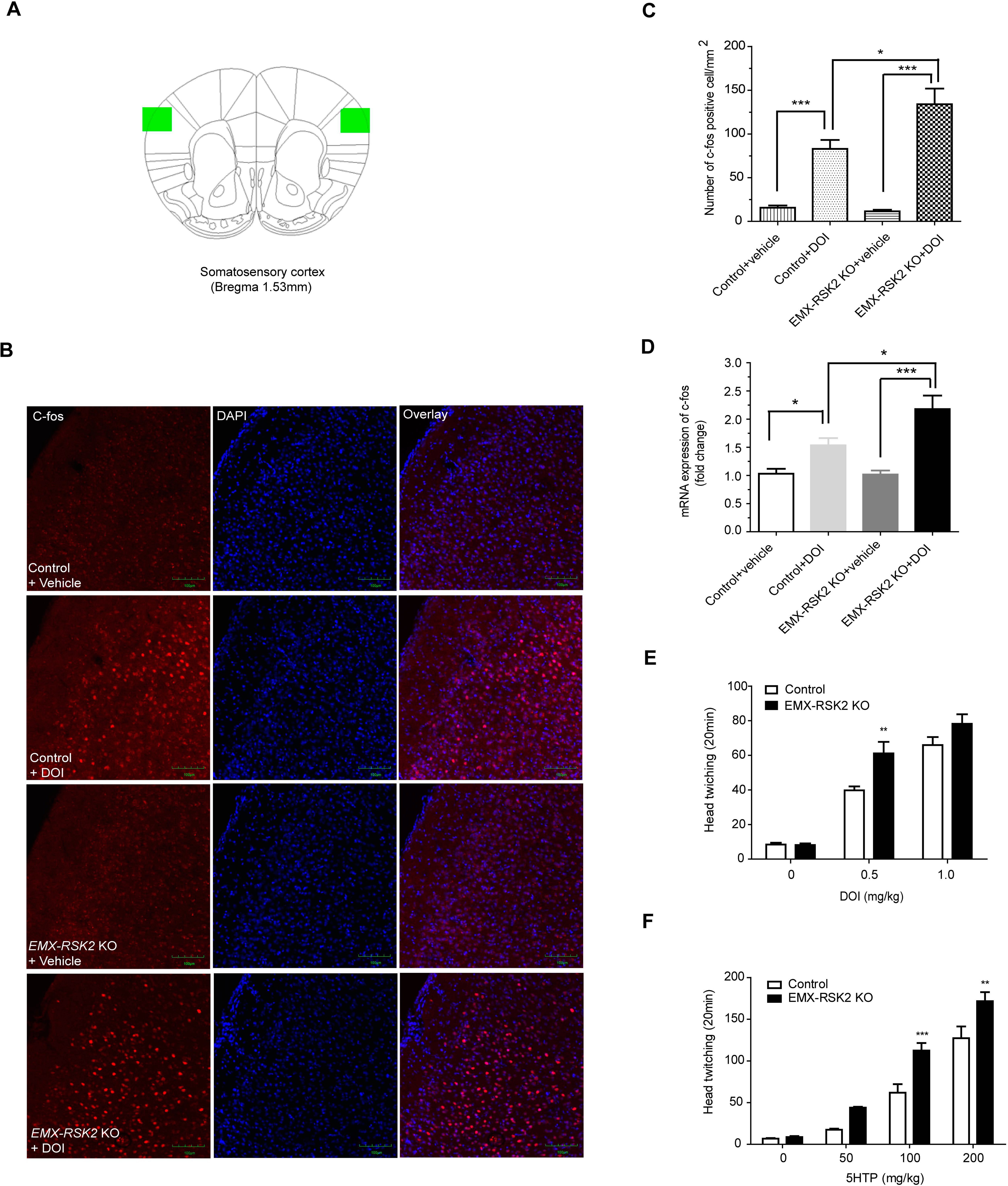
5-HT_2A_ receptor signaling is augmented in EMX-RSK2 KO mice. **A**. Schematic diagram of brain regions analyzed in DOI-induced c-fos activation experiment. Green boxes represent the region of somatosensory cortex analyzed in this study. Numbers indicate the distance from bregma in millimeters. **B, C**. DOI-induced c-fos activation is increased in EMX-RSK2 KO mice. Animals were injected with vehicle or 0.5 mg/kg DOI, 2 h later mice were perfused with 4% PFA, 40um frozen sections were cut and c-fos positive cells were determined by immunofluorescence (IF) staining. **B**. Representative figures of IF staining of DOI induced c-fos activation (green). Nuclei were counterstained with DAPI (blue). Sale bar, 100um. **C**. DOI-induced c-fos gene activation in somatosensory cortex is significantly increased in EMX-RSK2 KO mice compared to littermate control mice (F(1,24)=5.993, p=0.022, two-way ANOVA; n=7). **D**. DOI-induced c-fos expression is increased in EMX-RSK2 KO mice. Animals were injected with 0.5 mg/kg DOI, 1 h later frontal cortexes were dissected, mRNAs were extracted and transcript levels of c-fos were determined by real-time PCR. DOI-induced c-fos gene expression in frontal cortex was significantly increased in EMX-RSK2 KO mice (F(1,32)=4.93, p=0.0277 two-way ANOVA; n=9). **E**. HTR to the 5-HT_2A_ receptor partial agonist DOI was increased in EMX-RSK2 KO mice. Animals were injected with 0, 0.5, 1 mg/kg DOI and head twitches were scored over a 20 min period. DOI-induced head twitches were significantly increased in EMX-RSK2 KO mice compared with littermate control mice (F(2,36)=3.613, p=0.0372, two-way ANOVA; n=10). **F**. HTR to the 5-HT_2A_ receptor full agonist 5-HTP was increased in EMX-RSK2 KO mice. Animals were injected with 0, 50, 100, 200mg/kg 5-HTP and head twitches were scored over a 20 min period. DOI-induced head twitches were significantly increased in EMX-RSK2 KO mice compared with littermate control mice (F(3,24)=4.582, p=0.0113, twoway ANOVA; n=5). * p<0.05, ** p<0.01, *** p<0.001.

Next, we tested whether increased 5-HT_2A_ function in EMX-RSK2 KO mice leads to an increased head twitch response (HTR) in animals. The HTR is a reliable and extensively validated behavioral readout for 5-HT_2A_ agonism ( J. A. Allen et al., 2011) and has been shown to be elevated in PKCγ KO mice (B. J. Bowers et al., 2006). Following DOI (0, 0.5, 1.0 mg/kg) administration, both genotypes showed a dose-dependent increase in HTR over a 20 min period (Figure 5E). Notably, the HTR was significantly greater in EMX-RKS2 KO mice compared to littermate control mice (interaction of genotype and dose, F(2,36)=3.613, p=0.0372, effect of genotype, F(1,36)=9.827, p=0.0057, two-way ANOVA) (Figure 5E). Since DOI is a 5-HT_2A_ receptor partial agonist, we also treated mice with the full agonist 5-hydroxytryptophan (5-HTP) to fully activate this signaling pathway. Following 5-HTP (0, 50, 100, 200 mg/kg) administration, the HTR was increased in both genotypes (Figure 5F). Similar to DOI, we observed that 5-HTP (100 and 200 mg/kg) elicited a significantly greater HTR in EMX-RSK2 KO mice compared to the control mice (interaction of genotype and dose, F(3,24)=4.582, p=0.0113, effect of genotype, F(1,24)=19.57, p=0.0022, two-way ANOVA) (Figure 5F). Taken together, these results demonstrate that forebrain-specific RKS2 disruption potentiates 5-HT_2A_ function *in vivo*.

### 3.5 EMX-RSK2 KO mice show no alteration in 5-HT_2A_ receptor expression

We next quantified 5-HT2A receptor expression via receptor autoradiography (^125^I-DOI binding) and immunofluorescence microscopy to determine if altered receptor expression could account for the increased signaling in EMX-RSK2 KO mice. As shown in Figure 6A for both genotypes, we observed a high density of^125^I-DOI labeling in layer V of the cerebral cortex which is known to contain high levels of 5-HT_2A_ receptors (A. Pazos et al., 1985; J. F. Lopez-Gimenez et al., 1997). Quantitation revealed no significant difference between genotypes (t(8)=0.2448, p=0.8128, unpaired t-test) (Figure 6C) and complete displacement by ritanserin–a 5-HT_2A_/5-HT_2C_ selective antagonist–confirmed the specificity of^125^I-DOI binding. The expression of 5-HT_2A_ receptors in frontal cortex was further confirmed by immunostaining with a selective and previously validated 5-HT_2A_ receptor antibody (P. N. Yadav et al., 2011). We observed an overall diffuse laminar staining in the cerebral cortex and characteristic intensive band staining in the apical dendritic trunk and secondary branches of layer V pyramidal cells (Figure 6B). In agreement with the ^125^I-DOI labeling, no significant difference was observed in the intensity of layer V and peripheral layer staining between EMX-RSK2 KO mice and littermate control mice (F(1,8)=1.147, p=0.3153, two-way ANOVA) (Figure 6D). Taken together, our results demonstrate that augmented 5-HT_2A_ receptor signaling in EMX-RSK2 KO mice was not due to altered 5-HT_2A_ receptor expression in the forebrain.

**Figure 6.**
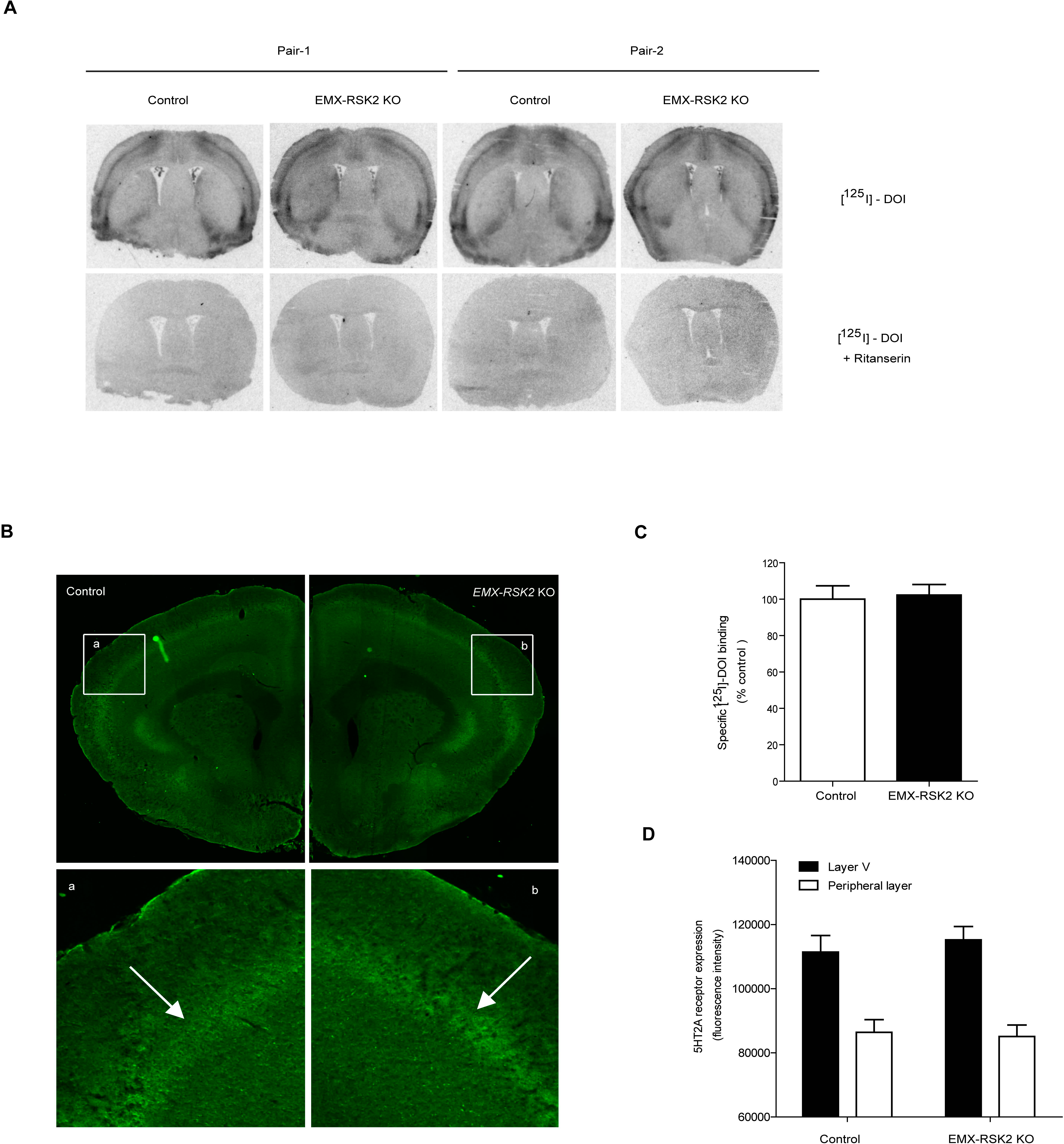
5-HT_2A_ receptor expression was not altered in EMX-RSK2 KO mice. **A**. Representative sections of 5-HT_2A_ receptor binding sites in the cerebral cortex of EMX-RSK2 KO mice and littermate control mice. Animal brains were cryosectioned and processed for ^125^I-DOI (200 pM) in vitro binding with/ without the presence of 1uM ritanserin. A clear signal in the layer V of the cerebral cortex was observed and the mean pixel intensity/area determined (upper panel). The specificity of ^125^I-DOI binding was shown that no signal was detected in the presence of ritanserin (lower panel). **B**. Representative sections of 5-HT_2A_ receptor immunoreactivity in cerebral cortex of EMX-RSK2 KO mice and littermate control mice. Animal brains were cryosectioned and stained with anti-5-HT_2A_ receptor antibody and Alexa 488 conjugated anti-rabbit secondary antibody. A clear layer V band (green) was detected in the cerebral cortex. Arrow, signals of 5-HT_2A_ receptor immunoreactivity in layer V. **C**. No difference in cortical ^125^I-DOI autoradiography tracing intensities was observed between genotypes in anatomically matched frontal cortex sections or examined (t(8)=0.2448, p=0.8128, unpaired t-test; n=5). **D**. No difference in cortical 5-HT_2A_ receptor abundance in layer V and peripheral layers was observed between genotypes in anatomically matched frontal cortex sections examined (F(1,8)=1.147, p=0.3153, two-way ANOVA; n=5).

## 4.0 DISCUSSION

The main findings of this paper are that (1) RSK2 exerts a tonic brake on 5-HT_2A_ signaling *in vivo,* and (2) that genetic deletion of RSK2 augments some of the psychotomimetic actions of amphetamine and PCP. Importantly, for these studies we generated a conditional RSK2 KO mouse to demonstrate that these behavioral effects can be attributed to selective deletion of RSK2 from the forebrain. These results provide compelling support for the hypothesis that RSK2 modulates 5-HT_2A_ signaling *in vivo,* dysregulation of which augments psychotomimetic drug actions.

As part of a large and ongoing effort to identify novel regulators of 5-HT_2A_ receptor signaling pathways, we identified RSK2 as a novel interacting protein that directly phosphorylates Ser-314 in the receptor third intracellular loop to decrease its signaling (D. J. Sheffler et al., 2006) (R. T. Strachan et al., 2009). Ultimately, it was revealed that RSK2 differentially regulates 5-HT_2A_ agonist signaling (R. T. Strachan et al., 2009; R. T. Strachan et al., 2010b) and that RSK2 is required for receptor tyrosine kinase (RTK)-mediated desensitization of 5-HT_2A_ receptors (R.T Strachan et al. 2010 Biochemistry). Consistent with these findings in cultured cells, we provide the first compelling evidence that RSK2 modulates 5-HT_2A_ signaling *in vivo*. First, genetic deletion of RSK2 leads to an augmentation of 5-HT_2A_ receptor signaling without altering apparent receptor expression. Second, genetic deletion of RSK2 increases psychostimulant-driven psychotomimetic responses (i.e., PCP- or AMPH-induced hyperlocomotion and disruption of PPI) that are incompletely blocked by the 5-HT_2A_-specific antagonist MDL100907, which we attribute to increased responsiveness of 5-HT_2A_ receptors (discussed below). However, it is possible that mechanisms outside of the 5-HT_2A_ receptor account for this MDL-insensitivity. For instance, RSK2 deletion could alter the activity of additional receptor systems that augment psychostimulant responses given that RSK2 regulates the activity of other GPCRs (P2Y-purinergic, PAR-1-thrombinergic, β1-adrenergic, and bradykinin-B receptors) (D. J. Sheffler et al., 2006). Moreover, Pereira et al. have shown dopaminergic dysregulation in the global RSK2 knockout mouse model (P. Marques Pereira et al., 2008). It is also possible that defects in neuronal growth due to impaired RSK2-dependent PLD1 activity contribute to the increased responsiveness of EMX-RSK2 KO mice to psychostimulants (M. R. Ammar et al., 2013), although the behavioral similarities between WT mice and forebrain deleted RSK2 mice argue against this possibility. More generally, our observation that 5-HT_2A_ receptor dysregulation is pro-psychotic and renders antagonists less effective is consistent with other *in vivo* studies in which 5-HT2A receptor interacting proteins were deleted (e.g., Cav1 and PSD-95 KO mice) (A.I. Abbas et al., 2009; J. A. Allen et al., 2011).

In the present study, we used multiple pharmacological approaches to evaluate 5-HT_2A_ receptor signaling in RSK2 mutant mice. In the first approach, we used the 5-HT_2A_ receptor selective antagonist MDL 100907 to block PCP- and AMPH-induced locomotion, which is typically used to evaluate 5-HT_2A_ signaling *in vivo*. In a second approach, we used the potent hallucinogen DOI to activate 5-HT_2A_ receptors in c-fos and HTR studies. Although DOI can activate both 5-HT_2A_ and 5-HT_2C_ receptors *in vitro,* several lines of evidence support the role of 5-HT_2A_ receptors, and not 5-HT_2C_ receptors, in mediating the activation of c-fos in the cortex and behavioral responses in animals. First, c-fos is primarily induced in cortical neurons expressing 5-HT_2A_ receptors (J. Gonzalez-Maeso et al., 2007). Second, pretreatment with the selective 5-HT_2A_ antagonist MDL 100,907 completely blocks DOI-induced c-fos expression, whereas pretreatment with the 5-HT_2C_ antagonist SB 206553 fails to reduce DOI-induced c-fos expression (J. L. Scruggs et al., 2000). Third, DOI and other hallucinogens do not induce c-fos expression and HTR behavior in 5-HT_2A_ KO mice (Gonzales-2007; Keiser et al, 2007), and Emx-Cre mediated restoration of 5-HT_2A_ receptor expression is sufficient to rescue the signaling and behavioral response (J. Gonzalez-Maeso et al., 2007). The finding that RSK2 mutant mice are more sensitive to the psychotomimetic effects of PCP is interesting in light of the observation of aschizophrenia-like psychosis seen in some CLS patients.

In summary, our *in vivo* data are consistent with the hypothesis that RSK2 tonically inhibits 5-HT_2A_ receptor signaling In this paradigm, genetic disruption of RSK2 augments 5-HT_2A_ receptor signaling and associated psychotic behaviors, suggesting that augmented 5-HT_2A_ receptor signaling is pro-psychotic and may contribute to the schizophrenia-like psychosis in some CLS patients. More generally, given the demonstrated clinical efficacy of antagonists with high affinity for 5-HT_2A_ receptors (B. L. Roth and Z. Xia, 2004; J. Besnard et al., 2012; H. Y. Meltzer and B. L. Roth, 2013), our findings suggest that RSK2 mutations, or mutations in other regulatory proteins, could alter the therapeutic efficacy of antipsychotic drugs in the clinic.

## 7.0 Acknowledgements

This work was supported by RO1MH61887 and U19MH82441 to BLR

## Footnotes

1 Abbreviations: RSK2: growth factor-regulated serine/threonine kinase, ribosomal S6 kinase 2; MAPK: multifunctional effector of the mitogen-activated protein kinase; CLS: Coffin-Lowry Sydnrome.

